# Prediction of Alzheimer’s Disease from Single Cell Transcriptomics Using Deep Learning

**DOI:** 10.1101/2023.07.07.548171

**Authors:** Aman Srivastava, Anjali Dhall, Sumeet Patiyal, Akanksha Arora, Akanksha Jarwal, Gajendra P. S. Raghava

## Abstract

Alzheimer’s disease (AD) is a progressive neurological disorder characterized by brain cell death, brain atrophy, and cognitive decline. Early diagnosis of AD remains a significant challenge in effectively managing this debilitating disease. In this study, we aimed to harness the potential of single-cell transcriptomics data from 12 Alzheimer’s patients and 9 normal controls (NC) to develop a predictive model for identifying AD patients. The dataset comprised gene expression profiles of 33,538 genes across 169,469 cells, with 90,713 cells belonging to AD patients and 78,783 cells belonging to NC individuals. Employing machine learning and deep learning techniques, we developed prediction models. Initially, we performed data processing to identify genes expressed in most cells. These genes were then ranked based on their ability to classify AD and NC groups. Subsequently, two sets of genes, consisting of 35 and 100 genes, respectively, were used to develop machine learning-based models. Although these models demonstrated high performance on the training dataset, their performance on the validation/independent dataset was notably poor, indicating potential overoptimization. To address this challenge, we developed a deep learning method utilizing dropout regularization technique. Our deep learning approach achieved an AUC of 0.75 and 0.84 on the validation dataset using the sets of 35 and 100 genes, respectively. Furthermore, we conducted gene ontology enrichment analysis on the selected genes to elucidate their biological roles and gain insights into the underlying mechanisms of Alzheimer’s disease. While this study presents a prototype method for predicting AD using single-cell genomics data, it is important to note that the limited size of the dataset represents a major limitation. To facilitate the scientific community, we have created a website to provide with code and service. It is freely available at https://webs.iiitd.edu.in/raghava/alzscpred.

**Key Points:**

- Predictive Model for Alzheimer’s Disease Using Single Cell Transcriptomics Data
- Overoptimization of models trained on single-cell genomics data.
- Application of dropout regularization technique of ANN for reducing overoptimization
- Ranking of genes based on their ability to predict patients’ Alzheimer’s Disease
- Standalone software package for predicting Alzheimer’s Disease

**Author’s Biography:**

1. Aman Srivastava is pursuing M. Tech. in Computational Biology from Department of Computational Biology, Indraprastha Institute of Information Technology, New Delhi, India.
2. Anjali Dhall is currently working as Ph.D. in Computational Biology from Department of Computational Biology, Indraprastha Institute of Information Technology, New Delhi, India.
3. Sumeet Patiyal is currently working as Ph.D. in Computational Biology from Department of Computational Biology, Indraprastha Institute of Information Technology, New Delhi, India.
4. Akanksha Arora is currently working as Ph.D. in Computational Biology from Department of Computational Biology, Indraprastha Institute of Information Technology, New Delhi, India.
5. Akanksha Jarwal is pursuing M. Tech. in Computational Biology from Department of Computational Biology, Indraprastha Institute of Information Technology, New Delhi, India.
6. Gajendra P. S. Raghava is currently working as Professor and Head of Department of Computational Biology, Indraprastha Institute of Information Technology, New Delhi, India.

## Introduction

Alzheimer’s disease (AD) poses a significant challenge to the medical industry and is a leading cause of dementia [1]. Globally, approximately 6.2 million individuals suffer from AD-induced dementia, and this number is projected to double every 20 years if a cure is not found [2,3]. AD is a progressive neurological disorder characterized by a decline in social, behavioral, and cognitive abilities, ultimately impairing independent functioning [4]. The presence of neuroinflammation, amyloid-beta peptide deposition, and tau neurofibrillary tangles are recognized as pathological biomarkers in AD [5]. However, it is important to note that their presence does not necessarily indicate the occurrence of AD-specific dementia, and the precise relationship to neurodegeneration remains unknown [6,7]. It is worth noting that although AD typically manifests in later stages of life, the pathogenesis often initiates much earlier than the onset of symptoms. Despite numerous studies and clinical trials focused on understanding and treating these pathological changes, no effective cure for AD currently exists [6]. Present treatment strategies primarily aim to alleviate symptoms and delay disease progression [8].

One of the reasons for the absence of the successful treatment methods for AD is the lack of knowledge of molecular underpinnings of cell-type specific responses for the pathogenesis of atrophy and neurodegeneration [9]. The bulk-tissue-level analysis may obscure the complexity of modifications between and within cells, particularly for rare cell types [10]. It is one of the significant challenges to understanding the complex human brain, which comprises a huge variety of cells that include billions of neurons of different subtypes [11]. Single-cell RNA sequencing provides an approach to analyze thousands of individual cells and study the compositional as well as functional changes within the cells to gain a clear insight into the mechanism of AD [12]. Recently, several studies have been conducted to identify the diagnostic and prognostic biomarkers of AD. For instance, Xiong et al., reported that KIR3DL2, PPP2R2B, and QPCT are upregulated in peripheral B cells while FRAT2, WWC3, and SPG20 are downregulated, and this gene panel of biomarkers can be used to diagnose and predict AD [13]. In addition, Yu et al. conducted a bioinformatics analysis and identified serum-based biomarkers (NEBL, EPB41L2, FGD4, and MARCKS) for AD [14].

In our study, we utilized single-cell RNA sequencing data from the GEO dataset, which included 169,496 nuclei from the prefrontal cortex of normal human subjects and AD patients (GSE157827) [15]. Through various machine learning and data mining approaches, we aimed to identify crucial gene-based biomarkers capable of stratifying AD and normal samples. Furthermore, to facilitate the scientific community, we have provided an end-to-end Python package. The complete workflow of our study is depicted in Figure 1.

**Figure 1:**
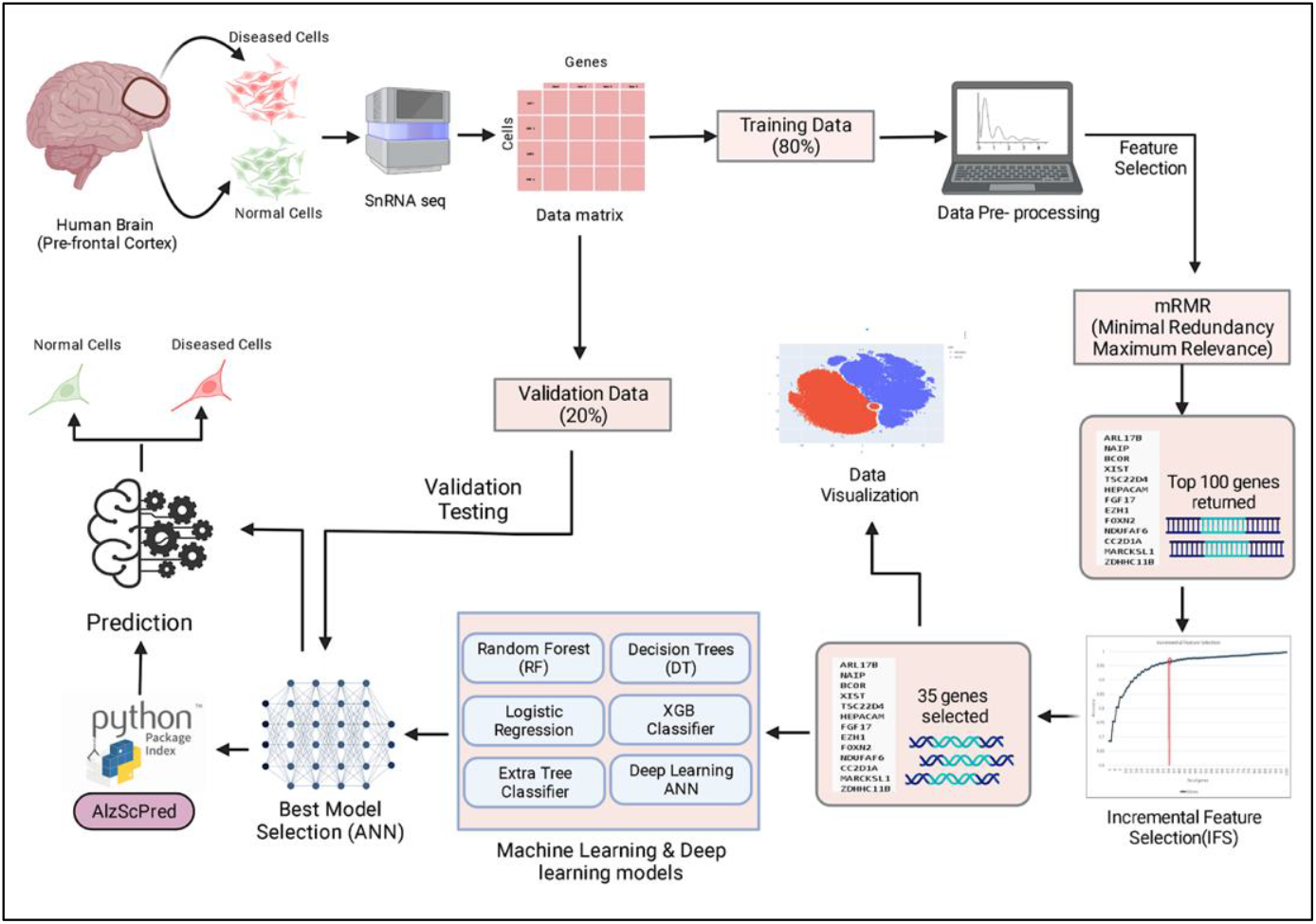
Workflow shows major steps involved in this study like data extraction, data processing, feature selection, and model development.

## Materials and methods

### Data Collection and Pre-processing

We accessed the single-cell gene expression profiles from the NCBI GEO repository, specifically the dataset with the GEO accession number GSE157827 [15]. The dataset comprises 21 samples, including 12 from individuals with Alzheimer’s disease (AD) and 9 from normal controls (NC). The gene expression profiles were generated using the Illumina NovaSeq 6000 sequencing platform, focusing on the prefrontal cortex of both AD and NC samples. The dataset consists of a total of 169,496 nuclei and encompasses 33,538 genes. We utilized the 10x single-cell sequencing data format, which included raw counts, barcodes, and gene files for each AD and NC sample. As the data obtained from the Cell Ranger pipeline was in sparse format, containing only non-zero entries to minimize file size, we transformed it into feature-barcode matrices using single-nucleus RNA sequencing (snRNA-seq). Subsequently, we removed outliers, including irrelevant genes and cells, using the scanty preprocessing library [16].

After the preprocessing steps, we identified 5,401 expressed genes, representing the number of genes with mapped reads in both AD and NC samples. Normalization was performed using the scanty.pp.normalize_total Python library. Additionally, we filtered out cells and genes containing NaN or zero values for each cell type to ensure data integrity. Finally, we divided the dataset into two parts: an 80% training dataset and a 20% validation dataset, enabling us to assess the performance of our models.

### Feature Selection Algorithm

In this study, we have used a feature selection approach in order to reduce the dimensions of the data matrix and to obtain an important set of genes that can accurately stratify the AD and NC samples. Here, we have used the information-theory-based feature-selection (IFA) method of minimum Redundancy and Maximum Relevance (mRMR), which has been previously implemented in similar types of studies analysis [17–19]. mRMR algorithm selects those genes/features which have a high correlation with the class (i.e., output) and a low correlation between themselves. The mRMR method not only takes into account the associations among the genes and samples but also reduces the redundancy within genes. For example, it selects only the most representative gene out of similar types of genes. Here, we have selected the top 100 most relevant genes and then applied an incremental feature selection algorithm to find out the optimal set of genes as biomarkers [20].

### Model Development

Various machine learning and deep learning techniques have been implemented in this study to develop the models for the classification of NC and AD samples. The machine learning algorithms include decision tree (DT), random forest (RF), logistic regression (LR), XGBoost (XGB), and extra tree classifiers (ET) [21–24]. The non-parametric supervised learning models serve as the foundation for the DT algorithm. The goal was to create a model by learning the decision rules from the data features that may predict the response variable. RF is an ensemble-based approach for classification that predicts the response variable using each individual tree while fitting a variety of DTs during training. Averaging DTs increases model control over overfitting and prediction accuracy. The logistic or logit model, which offers a class or event probability, can be obtained using the LR approach. To develop a model that can predict a class or response variable, the logistic function is used. This approach resembles multiple linear regression. However, the dependent variable, in this case, is binomial instead. Scalable tree boosting classification algorithm XGB makes a final prediction after several iterations. It makes use of the ensemble method, which combines a number of models to make the final forecast. We created models that can forecast the response variable or class of the sample by using parameters tuned on the training dataset.

ML models were observed to be performing well on the training dataset. However, the performance was not comparable when checked on an independent validation set. In order to overcome this problem, we have used deep learning-based models to stratify the samples, which use a customized artificial neural network (ANN). An ANN is composed of a network of interconnected systems or nodes known as artificial neurons, which are generally designed like neurons in the human brain [25]. Our neural network architecture consists of 1 input layer, 3 hidden layers, and 1 output layer with a dropout of 0.3 at each step to reduce the overfitting of the deep learning model. The final output is the predicted label which tells whether the sample is diseased or not. Scikit-Learn, TensorFlow, and Keras Python libraries were used to implement the machine learning and deep learning algorithms [26,27].

### Cross-validation technique

To ensure the robustness and reliability of our prediction models, we employed the 5-fold cross-validation technique in combination with external validation. This approach allows us to evaluate the models using both internal and external datasets. The 80% training dataset was utilized for internal validation, where the data was split into five equal sets or folds, ensuring an equal representation of positive and negative samples in each fold. Subsequently, the models were trained using four-folds, while the fifth fold served as the test set. This process was repeated five times, with each fold acting as the test set once, and the performance metrics were calculated for each iteration. Finally, the performance results from all five sets were averaged to obtain the final performance assessment. In addition to the internal validation, we performed external validation using the remaining 20% of the data. This independent dataset served as an external benchmark to assess the generalizability of our models. By evaluating the performance on both internal and external datasets, we obtained a comprehensive understanding of the predictive capabilities and reliability of our models. This technique has been previously used in many studies [28,29].

### Performance measures

We employed established evaluation metrics to assess the effectiveness of various prediction models. We employed both threshold-dependent and independent parameters in this study. We have used the following equations to quantify threshold-dependent parameters such as sensitivity (Sens), specificity (Spec), precision, F1-Score, and accuracy (Acc). In order to evaluate the effectiveness of the models, we additionally employed the typical threshold-independent parameter Area Under the Receiver Operating Characteristic (AUROC) curve.

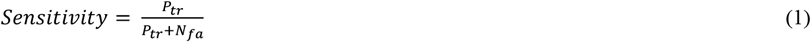

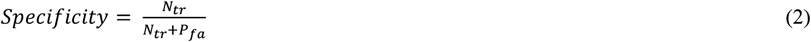

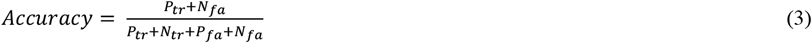

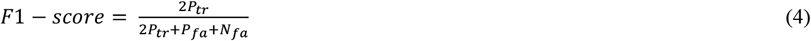

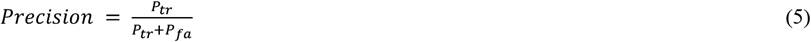

Where *P*_*tr*_ is truly positive, *N*_*tr*_ is truly negative, *P*_*fa*_ is false positive, and *N*_*fa*_ is false negative.

## Results

### Feature Selection

In this work, we have used snRNA-seq expression profiles of 12 AD and 9 NC samples incorporating 5401 genes and 128706 cells. We used the mRMR technique to shortlist a set of 100 genes that were highly relevant in regard to the detection of Alzheimer’s disease. The selected top 100 genes are provided in the supplementary table S1. To make our prediction model even more accurate, we wanted to obtain a smaller set of genes that play an important role in the detection of Alzheimer’s disease. To achieve the same, we employed the incremental feature selection (IFS) technique, where we plot an IFS curve for all the combinations of genes (features) on the x-axis and the accuracy value obtained from all sets on the y-axis as shown in Figure 2. The optimum accuracy was obtained when the model was trained with 35 biomarkers, as after that, there was no significant improvement in the accuracy of the model. A list of the set of 35 selected genes is given in supplementary table S2.

**Figure 2:**
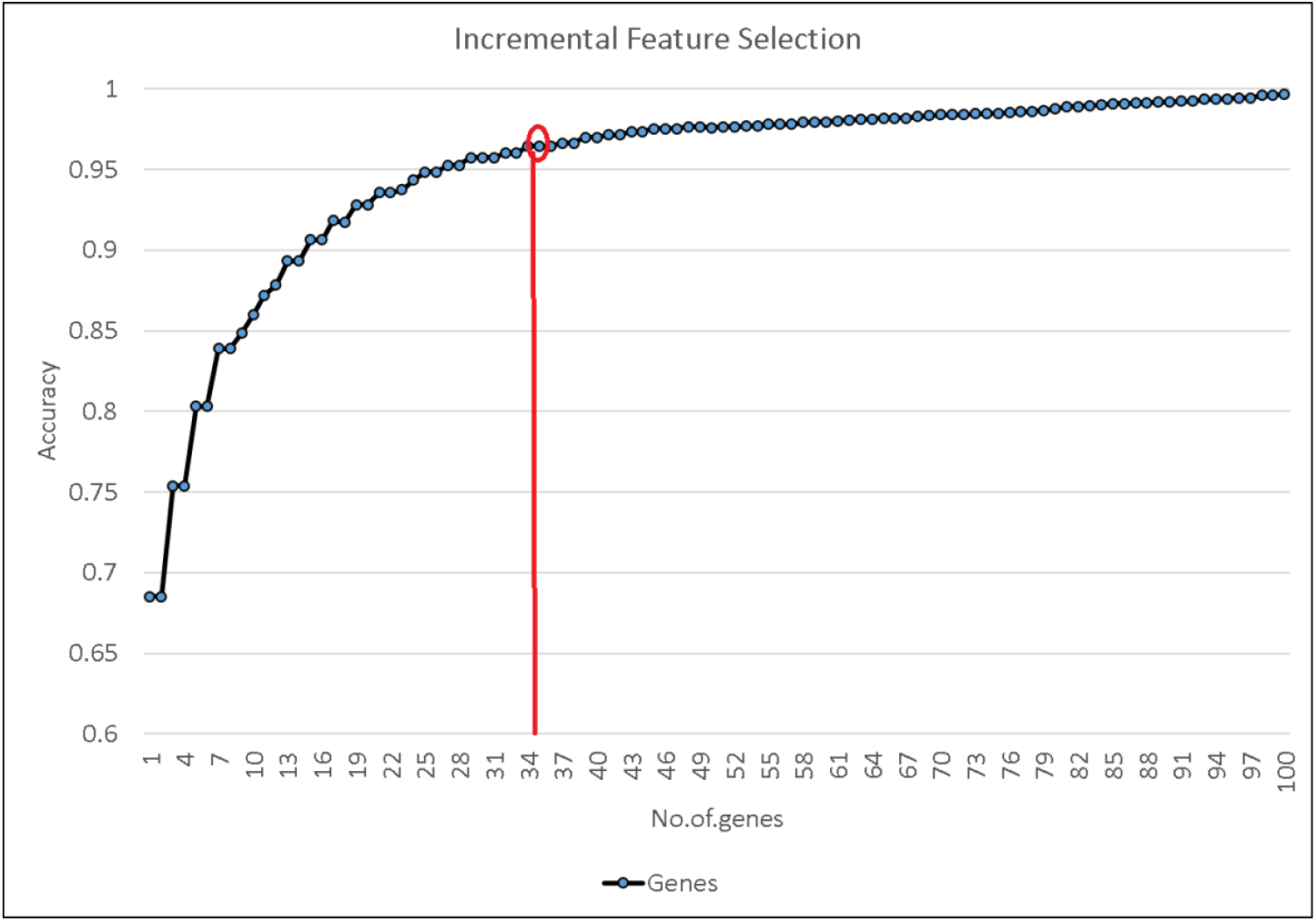
Accuracy of models developed using a set of features selected using the incremental feature selection (IFS) technique.

**Figure 2:**
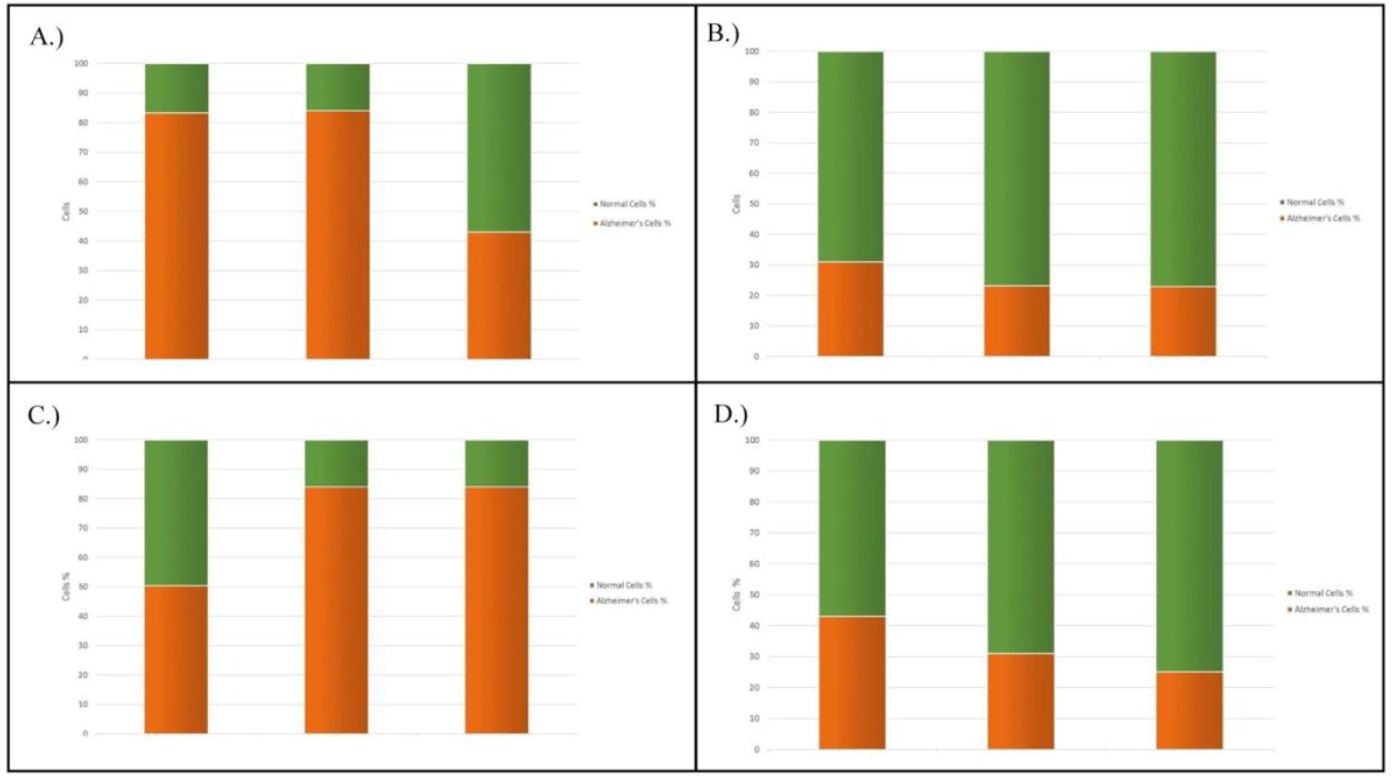
Plot for patient-wise analysis using single-cell RNA profile

### Performance of classification models

Initially, we applied various machine learning algorithms like DT, ET, RF, LR, and XGB, and we found that most of the models performed well on the training dataset, but the performance was not comparable on validation data (See Table 1). In order to improve the prediction performance, we have applied deep learning. It was observed that deep learning-based models outperform all the machine learning models on the validation dataset. As shown in Table 1, we were able to achieve a maximum AUROC of 0.84 and an accuracy of 82.00% on an independent validation dataset for a set of 100 genes selected using the mRMR feature selection technique. For a set of 35 genes that were shortlisted using the IFS technique, we obtained the maximum AUROC of 0.75 and an accuracy of 74.00% on the independent validation set. The results for the top 35 selected genes are given in Table 1.

**Table 1:** Performance of various machine learning and deep learning models using top 100 features (mRMR) and top 35 features selected using IFS.

| Top 100 genes |  |  |  |  |  |  |  |  |  |  |
| --- | --- | --- | --- | --- | --- | --- | --- | --- | --- | --- |
| Model | Training Dataset |  |  |  |  | Validation Dataset |  |  |  |  |
|  | Sens | Spec | Acc (%) | f1-Score | AUROC | Sens | Spec | Acc (%) | f1-Score | AUROC |
| **DT** | 0.99 | 1 | 100 | 0.99 | 0.99 | 0.97 | 0.02 | 40 | 0.04 | 0.49 |
| **RF** | 0.96 | 1 | 98 | 0.98 | 0.98 | 0.89 | 0.09 | 41 | 0.16 | 0.49 |
| **LR** | 0.99 | 1 | 100 | 0.99 | 0.99 | 0.68 | 0.18 | 38 | 0.28 | 0.43 |
| **XGB** | 0.99 | 1 | 99 | 0.99 | 0.99 | 0.36 | 0.55 | 47 | 0.43 | 0.45 |
| **ET** | 0.99 | 1 | 100 | 0.99 | 0.99 | 0.85 | 0.13 | 43 | 0.23 | 0.5 |
| **ANN** | 0.99 | 1 | 99 | 0.99 | 0.99 | 0.86 | 0.77 | 82 | 0.8 | 0.84 |
| Top 35 genes |  |  |  |  |  |  |  |  |  |  |
| Model | Training Dataset |  |  |  |  | Validation Dataset |  |  |  |  |
|  | Sens | Spec | Acc (%) | f1-Score | AUROC | Sens | Spec | Acc (%) | f1-Score | AUROC |
| **DT** | 1 | 0.93 | 96.39 | 0.97 | 0.97 | 0.92 | 0.07 | 42 | 0.13 | 0.5 |
| **RF** | 0.91 | 1 | 95.93 | 0.95 | 0.96 | 0.81 | 0.15 | 42 | 0.25 | 0.48 |
| **LR** | 1 | 0.93 | 96.39 | 0.97 | 0.97 | 0.62 | 0.4 | 38 | 0.38 | 0.38 |
| **XGB** | 1 | 0.93 | 96.38 | 0.97 | 0.97 | 0.54 | 0.41 | 46 | 0.47 | 0.48 |
| **ET** | 0.91 | 1 | 96 | 0.95 | 0.96 | 0.5 | 0.43 | 46 | 0.46 | 0.47 |
| **ANN** | 1 | 0.93 | 96.4 | 0.97 | 0.97 | 0.75 | 0.66 | 74 | 0.71 | 0.75 |

#DT: Decision Tree; RF: Random Forest; LR: Logistic Regression; XGB: XGBoost; ET: Extra Tree Classifiers; ANN: Artificial Neural Network; AUROC: Area Under the Receiver Operating Characteristics curve; Sens: Sensitivity, Spec: Specificity; Acc: Accuracy

### Patient wise analysis

To test our final model based on deep learning, we fed the data containing a full single-cell sequence profile for each patient and saw how accurately it predicted the disease in each subject. The results for the patient-wise analysis for the top 100 genes and the top 35 genes are given in Table 2 and Figure 3.

**Table 2:** Patient-wise analysis using single-cell RNA sequence profile

| Human Subjects | 100 Genes |  | 35 Genes |  |
| --- | --- | --- | --- | --- |
|  | Predicted | Predicted | Predicted | Predicted |
|  | Diseased Cells | Normal Cells | Diseased Cells | Normal Cells |
| **Alzheimer Test 1** | 43.00% | 57.00% | 50.41% | 49.58% |
| **Alzheimer Test 2** | 83.27% | 16.70% | 83.98% | 16.01% |
| **Alzheimer Test 3** | 83.97% | 16.02% | 83.97% | 16.02% |
| **Normal Test 1** | 31.00% | 69.00% | 43.00% | 57.00% |
| **Normal Test 2** | 23.21% | 76.80% | 31.00% | 69.00% |
| **Normal Test3** | 22.98% | 77.01% | 25.17% | 74.80% |

### Biological functions of selected genes

After feature selection using mRMR and IFS, about 35 genes were extracted, which could serve as potential diagnostic biomarkers of Alzheimer’s. We then performed Gene Ontology (GO) Enrichment Analysis on these extracted 35 genes to map the biological functions of the selected genes. The Go enrichment analysis results are shown in Table 3. Most of the Genes are involved in the Binding Activity and catalytic activity of various metabolic processes. Other activities associated with reported genes are ATP-dependent activity, molecular function regulator, transcription regulator activity, molecular transducer activity, and transporter activity.

**Table 3:** Gene Ontology (GO) enrichment analysis for the selected 35 genes

| GO term | Activity | Genes |
| --- | --- | --- |
| (GO:0140657) | ATP-dependent activity | CHD7 |
| (GO:0005488) | Binding | CHD7, ARL17B, UBE2Z, FGF17, OTUD7B, CDKL1, MARCKSL1, FOXN2,<br>NAIP, PER1, CC2D1A, AC090517, LPAR1, PWWP2A, CDK18 |
| (GO:0003824) | Catalytic Activity | ZDHHC11B, CHD7, UBE2Z, HIBADH, OTUD7B, |
|  |  | CDKL1, NAIP, CDK18, PLPPR2 |
| (GO:0098772) | Molecular Function<br>Regulator | UBE2Z, FGF17, NAIP |
| (GO:0060089) | Molecular Transducer<br>Activity | PLXNB1, FGF17, LPAR1 |
| (GO:0140110) | Transcription Regulator<br>Activity | BCOR, FOXN2, CC2D1A, AC090517 |
| (GO:0005215) | Transporter Activity | SLC25A13 |

## Discussions

Alzheimer’s has now been recognized in the category of world health concerns. It accounts for nearly 60-70% of cases of dementia [30]. There are several criteria that have been proposed for the screening and diagnosis of AD, including physical symptoms, bodily fluids, and imaging studies. Despite this, there is currently no cure for AD, and the only effective therapy involves the management of symptoms. Various drugs such as galantamine, donepezil, and rivastigmine are prescribed to increase memory power and brain alertness, but they cannot control the disease progression [31]. Numerous studies have demonstrated that altering lifestyle practices, including food and exercise, can improve brain health and lessen AD without needing medical attention [32]. Hence, it is recommended as a first-line prevention strategy for all AD patients. Significant neuronal loss and neuropathological abnormalities can harm several brain regions by the time the disease is identified [33]. To prevent this, the study aims to identify some specific biomarkers which could aid in early screening and diagnosis of Alzheimer’s disease from the Single Cell genome.

In this study, we have used various machine learning models and a deep learning model to classify NC cells and AD cells from their single-cell RNA-seq profile. Also, patient-wise analysis to classify the samples is also done in this study on a total of 21 samples. We compared the deep learning model with other traditional machine learning models to find out that the deep learning model performed well very against all the traditional machine learning models. Initially, the dataset was quite large, with a very high number of features (>30000). The data were then pre-processed, and the feature count was reduced to a significantly lower number (=5000). Since many features were correlated and redundant, so a feature selection method known as mRMR was applied to get a minimal set of features, which could be helpful in classifying the samples. Out of these 5000 features, mRMR was applied to extract the top 100 features with minimal redundancy and maximal relevance, which achieved an AUROC of 0.84 on an independent validation set using deep learning. The IFS method was then applied to select an even smaller gene set and obtain an optimal number of biomarkers. We found about 35 genes (features) to be optimal, and they were able to distinguish the AD patients from NC individuals with an accuracy of 74% and AUROC of 0.75 on the independent validation set. In addition, we performed Gene Ontology (GO) enrichment analysis in our study to gain insight into the mechanism and role of selected genes in the development of Alzheimer’s disease. Silencing of lncRNA X-inactive specific transcript (XIST) was found to be directly connected to an increase in Alzheimer’s symptoms. The XIST gene was found to be upregulated in AD models in both in vitro and in vivo studies [34]. Gene “TSC22D4” in a study with Japanese Alzheimer’s patients showed to be significantly differentially regulated as compared to normal controls [35]. “FGF17”, i.e., Fibroblast growth factor dysregulation, has been reported in various brain-related (neurological and psychiatric) disorders. It has been shown to be altered in epileptogenesis [36]. Transcriptional regulatory changes are prominent features of brain diseases, transcriptional changes in the gene “FOXN2” highlights the convergence of genetic risk with psychiatric and neurodegenerative disorders [37].

Genes such as “CDK18” have uncharacterized mechanisms by which they may promote AD neurodegeneration, and this increases the probability that their inhibition may cause protection against AD development pathology [38]. There are also genes such as ‘BCOR,’ ‘AC090517.4’, ‘LGI4’, ‘SLC25A13’, and ‘ZBED5’, which have not yet been reported in any connection with Alzheimer’s disease progression and development. Thus, further deep studies are required to evaluate these biomarkers, and these could act as novel findings. Genes ‘SCD’ and ‘UBE2Z’ have been reported in various studies related to cognitive impairment and diagnosis or treatment of Alzheimer’s disease [39,40]. About 21 genes out of the 35 genes reported in this study have been observed to have a connection to Alzheimer’s disease and a relation with neurodegeneration. Other findings need further study and research.

The limitation of this study is that we have only evaluated a total of 21 samples from both normal and Alzheimers affected patients. The data was obtained from the prefrontal cortex of the brain. More data from areas like the hippocampus and amygdala can be obtained to add to this dataset for identifying further promising biomarkers. In addition, more studies on the identified genes in clinical setups are required for in-depth analysis and their role in how they affect and cause Alzheimer’s disease progression.

## Funding Source

The current work has been supported by the Department of Biotechnology (DBT) grant BT/PR40158/BTIS/137/24/2021.

## Conflict of interest

The authors declare no competing financial and non-financial interests.

## Ethics Approval

Not applicable

## Consent to Participate

Not applicable

## Conflict of Publication

Not applicable

## Acknowledgements

Authors are thankful to the Department of Biotechnology (DBT grant BT/PR40158/BTIS/137/ 24/2021), Council of Scientific & Industrial Research (CSIR), and Department of Science and Technology (DST-INSPIRE) for fellowships and the financial support. Authors are also thankful to Department of Computational Biology, IIITD New Delhi for infrastructure and facilities.

## Authors’ contributions

AS and AD collected and processed the datasets. AS, AD and SP implemented the algorithms and developed the prediction models. AS, AD, SP and AJ analyzed the results. AS and AA created the back end of the web server and created the front-end user interface. AS, AD, SP, AA and AJ penned the manuscript. GPSR conceived and coordinated the project. All authors have read and approved the final manuscript.

## Notes

### Competing Interest Statement

The authors have declared no competing interest.

## References

[1] Mayeux, R., Stern, Y., Epidemiology of Alzheimer disease. Cold Spring Harb Perspect Med 2012, 2.

[2] Yiannopoulou, K.G., Papageorgiou, S.G., Current and Future Treatments in Alzheimer Disease: An Update. J Cent Nerv Syst Dis 2020, 12, 1179573520907397.

[3] Cummings, J., Lee, G., Nahed, P., Kambar, M.E.Z.N., et al., Alzheimer’s disease drug development pipeline: 2022. Alzheimers Dement (N Y) 2022, 8, e12295.

[4] DeTure, M.A., Dickson, D.W., The neuropathological diagnosis of Alzheimer’s disease. Mol Neurodegener 2019, 14, 32.

[5] Murphy, M.P., LeVine, H., Alzheimer’s disease and the amyloid-beta peptide. J Alzheimers Dis 2010, 19, 311–23.

[6] Kim, D., Kwon, H.J., Hyeon, T., Magnetite/Ceria Nanoparticle Assemblies for Extracorporeal Cleansing of Amyloid-β in Alzheimer’s Disease. Adv Mater 2019, 31, e1807965.

[7] Srivastava, S., Ahmad, R., Khare, S.K., Alzheimer’s disease and its treatment by different approaches: A review. Eur J Med Chem 2021, 216, 113320.

[8] Yiannopoulou, K.G., Papageorgiou, S.G., Current and Future Treatments in Alzheimer Disease: An Update. J Cent Nerv Syst Dis 2020, 12, 1179573520907397.

[9] Mathys, H., Davila-Velderrain, J., Peng, Z., Gao, F., et al., Single-cell transcriptomic analysis of Alzheimer’s disease. Nature 2019, 570, 332–337.

[10] De Strooper, B., Karran, E., The Cellular Phase of Alzheimer’s Disease. Cell 2016, 164, 603–15.

[11] Silbereis, J.C., Pochareddy, S., Zhu, Y., Li, M., Sestan, N., The Cellular and Molecular Landscapes of the Developing Human Central Nervous System. Neuron 2016, 89, 248–68.

[12] Hwang, B., Lee, J.H., Bang, D., Single-cell RNA sequencing technologies and bioinformatics pipelines. Exp Mol Med 2018, 50, 1–14.

[13] Gao, W., Xiong, Y., Li, Q., Yang, H., Inhibition of toll-like receptor signaling as a promising therapy for inflammatory diseases: A journey from molecular to nanotherapeutics. Front Physiol 2017, 8.

[14] Yu, Q.-S., Feng, W.-Q., Shi, L.-L., Niu, R.-Z., Liu, J., Integrated Analysis of Cortex Single-Cell Transcriptome and Serum Proteome Reveals the Novel Biomarkers in Alzheimer’s Disease. Brain Sci 2022, 12.

[15] Lau, S.-F., Cao, H., Fu, A.K.Y., Ip, N.Y., Single-nucleus transcriptome analysis reveals dysregulation of angiogenic endothelial cells and neuroprotective glia in Alzheimer’s disease. Proc Natl Acad Sci U S A 2020, 117, 25800–25809.

[16] Wolf, F.A., Angerer, P., Theis, F.J., SCANPY: large-scale single-cell gene expression data analysis. Genome Biol 2018, 19, 15.

[17] Zulfiqar, H., Ahmed, Z., Kissanga Grace-Mercure, B., Hassan, F., et al., Computational prediction of promotors in Agrobacterium tumefaciens strain C58 by using the machine learning technique. Front Microbiol 2023, 14, 1170785.

[18] Arabi, M., Nazari, M., Salahshour, A., Jenabi, E., et al., A machine learning-based economics for prediction of thyroid nodule malignancies. Endocrine 2023.

[19] Kaplan, E., Altunisik, E., Ekmekyapar Firat, Y., Datta Barua, P., et al., Novel nested patch-based feature extraction model for automated Parkinson’s Disease symptom classification using MRI images. Comput Methods Programs Biomed 2022, 224, 107030.

[20] Cheng, Q., Li, J., Fan, F., Cao, H., et al., Identification and Analysis of Glioblastoma Biomarkers Based on Single Cell Sequencing. Front Bioeng Biotechnol 2020, 8, 167.

[21] Bulac, C., Bulac, A., in: Advanced Solutions in Power Systems: HVDC, FACTS, and AI Techniques, 2016.

[22] Breiman, L., Random forests. Mach Learn 2001.

[23] Stoltzfus, J.C., Logistic regression: a brief primer. Acad Emerg Med 2011, 18, 1099–104.

[24] Chen, T., Guestrin, C., in: Proceedings of the ACM SIGKDD International Conference on Knowledge Discovery and Data Mining, 2016.

[25] Wang, S.-C., in: Interdisciplinary Computing in Java Programming, Springer US, Boston, MA 2003, pp. 81–100.

[26] Pedregosa, F., Varoquaux, G., Gramfort, A., Michel, V., et al., Scikit-learn: Machine Learning in Python 2012.

[27] Abadi, M., Barham, P., Chen, J., Chen, Z., et al., TensorFlow: A system for large-scale machine learning 2016.

[28] Aggarwal, S., Dhall, A., Patiyal, S., Choudhury, S., et al., An ensemble method for prediction of phage-based therapy against bacterial infections. Front Microbiol 2023, 14, 1148579.

[29] Arora, A., Patiyal, S., Sharma, N., Devi, N.L., et al., A random forest model for predicting exosomal proteins using evolutionary information and motifs. bioRxiv 2023, 2023.01.30.526378.

[30] Chanda, K., Mukhopadhyay, D., LncRNA Xist, X-chromosome Instability and Alzheimer’s Disease. Curr Alzheimer Res 2020, 17, 499–507.

[31] Breijyeh, Z., Karaman, R., Comprehensive Review on Alzheimer’s Disease: Causes and Treatment. Molecules 2020, 25.

[32] Bhatti, G.K., Reddy, A.P., Reddy, P.H., Bhatti, J.S., Lifestyle Modifications and Nutritional Interventions in Aging-Associated Cognitive Decline and Alzheimer’s Disease. Front Aging Neurosci 2019, 11, 369.

[33] Serrano-Pozo, A., Frosch, M.P., Masliah, E., Hyman, B.T., Neuropathological alterations in Alzheimer’s disease. Cold Spring Harb Perspect Med 2011, 1, a006189.

[34] Yue, D., Guanqun, G., Jingxin, L., Sen, S., et al., Silencing of long noncoding RNA XIST attenuated Alzheimer’s disease-related BACE1 alteration through miR-124. Cell Biol Int 2020, 44, 630–636.

[35] Humphries, C., Kohli, M.A., Whitehead, P., Mash, D.C., et al., Alzheimer disease (AD) specific transcription, DNA methylation and splicing in twenty AD-associated loci. Mol Cell Neurosci 2015, 67, 37–45.

[36] Turner, C.A., Eren-Koçak, E., Inui, E.G., Watson, S.J., Akil, H., Dysregulated fibroblast growth factor (FGF) signaling in neurological and psychiatric disorders. Semin Cell Dev Biol 2016, 53, 136–43.

[37] Pearl, J.R., Colantuoni, C., Bergey, D.E., Funk, C.C., et al., Genome-Scale Transcriptional Regulatory Network Models of Psychiatric and Neurodegenerative Disorders. Cell Syst 2019, 8, 122–135.e7.

[38] Chaput, D., Kirouac, L., Stevens, S.M., Padmanabhan, J., Potential role of PCTAIRE-2, PCTAIRE-3 and P-Histone H4 in amyloid precursor protein-dependent Alzheimer pathology. Oncotarget 2016, 7, 8481–97.

[39] Hamilton, L.K., Moquin-Beaudry, G., Mangahas, C.L., Pratesi, F., et al., Stearoyl-CoA Desaturase inhibition reverses immune, synaptic and cognitive impairments in an Alzheimer’s disease mouse model. Nat Commun 2022, 13, 2061.

[40] Lim, K.-H., Joo, J.-Y., Predictive Potential of Circulating Ube2h mRNA as an E2 Ubiquitin-Conjugating Enzyme for Diagnosis or Treatment of Alzheimer’s Disease. Int J Mol Sci 2020, 21.

