## Supplementary Table S1 and Supplementary Table S2 for "Prediction of Alzheimer’s Disease from Single Cell Transcriptomics Using Deep Learning"

**Mailing Address of Authors**

Aman Srivastava (AS):

Anjali Dhall (AD):

Sumeet Patiyal (SP):

Akanksha Arora (AA):

Akanksha Jarwal (AJ):

Gajendra P. S. Raghava (GPSR):

***Corresponding Author**

Prof. Gajendra P. S. Raghava

Head and Professor

Department of Computational Biology

Indraprastha Institute of Information Technology, Delhi

Okhla Industrial Estate, Phase III, (Near Govind Puri Metro Station)

New Delhi, India – 110020

Office: A-302 (R&D Block)

Website: http://webs.iiitd.edu.in/raghava/

**Supplementary Table S1: Top 100 genes obtained by mRMR algorithm**

| Rank | Genes | Rank | Genes | Rank | Genes | Rank | Genes |
|---|---|---|---|---|---|---|---|
| 01 | ARL17B | 26 | HIBADH | 51 | DDX3X | 76 | APOD |
| 02 | NAIP | 27 | ZBED5 | 52 | NSL1 | 77 | KIF9-AS1 |
| 03 | BCOR | 28 | PTDSS2 | 53 | TMED10 | 78 | TYW1 |
| 04 | XIST | 29 | ATG4B | 54 | UGT8 | 79 | GNAI2 |
| 05 | TSC22D4 | 30 | PWWP2A | 55 | SNX1 | 80 | BAZ1B |
| 06 | HEPACAM | 31 | XRRA1 | 56 | ALG13 | 81 | MBTPS1 |
| 07 | FGF17 | 32 | OTUD7B | 57 | LINC00320 | 82 | CDH4 |
| 08 | EZH1 | 33 | SCD | 58 | RAD9A | 83 | RAB40B |
| 09 | FOXN2 | 34 | UBE2Z | 59 | RGS12 | 84 | SPP1 |
| 10 | NDUFAF6 | 35 | PIGQ | 60 | ST13 | 85 | GPBP1L1 |
| 11 | CC2D1A | 36 | PCMTD2 | 61 | PTN | 86 | FSCN1 |
| 12 | MARCKSL1 | 37 | COL4A5 | 62 | USP8 | 87 | CAPZA1 |
| 13 | ZDHHC11B | 38 | ARFIP1 | 63 | EDF1 | 88 | SPPL2B |
| 14 | PLXNB1 | 39 | CCND3 | 64 | SLCO1A2 | 89 | MED15 |
| 15 | PLPPR2 | 40 | FOXK2 | 65 | NUP153 | 90 | C1GALT1 |
| 16 | AC090517.4 | 41 | CPOX | 66 | SYNRG | 91 | ITGB3BP |
| 17 | CDK18 | 42 | STXBP3 | 67 | EIF3E | 92 | CCDC82 |
| 18 | LGI4 | 43 | ITPKB | 68 | LPCAT4 | 93 | CDK12 |
| 19 | CHD7 | 44 | TBCB | 69 | ARMCX4 | 94 | EGLN1 |
| 20 | RBMX | 45 | SRSF10 | 70 | PREX2 | 95 | CCDC57 |
| 21 | CDKL1 | 46 | SPTLC2 | 71 | APC2 | 96 | SEL1L |
| 22 | DNAJC7 | 47 | LYPLAL1 | 72 | SLC38A9 | 97 | CHORDC1 |
| 23 | SLC25A13 | 48 | FAM107B | 73 | UTP23 | 98 | ATG10 |
| 24 | PER1 | 49 | PDIA2 | 74 | RCN2 | 99 | C4orf48 |
| 25 | LPAR1 | 50 | C1orf61 | 75 | PRR14L | 100 | AC097103.2 |

**Supplementary Table S2: Top 35 gene subset obtained using IFS**

| Rank | Gene | Rank | Gene | Rank | Gene |
|---|---|---|---|---|---|
| 01 | ARL17B | 13 | ZDHHC11B | 25 | LPAR1 |
| 02 | NAIP | 14 | PLXNB1 | 26 | HIBADH |
| 03 | BCOR | 15 | PLPPR2 | 27 | ZBED5 |
| 04 | XIST | 16 | AC090517.4 | 28 | PTDSS2 |
| 05 | TSC22D4 | 17 | CDK18 | 29 | ATG4B |
| 06 | HEPACAM | 18 | LGI4 | 30 | PWWP2A |
| 07 | FGF17 | 19 | CHD7 | 31 | XRRA1 |
| 08 | EZH1 | 20 | RBMX | 32 | OTUD7B |
| 09 | FOXN2 | 21 | CDKL1 | 33 | SCD |
| 10 | NDUFAF6 | 22 | DNAJC7 | 34 | UBE2Z |
| 11 | CC2D1A | 23 | SLC25A13 | 35 | PIGQ |
| 12 | MARCKSL1 | 24 | PER1 | | |
